# *Cse1l* Regulates Neural Crest Cell Survival and is Critical for Craniofacial and Cardiac Development

**DOI:** 10.1101/2025.05.06.652282

**Authors:** Paul P. R. Iyyanar, Rolf W. Stottmann

**Affiliations:** Steve and Cindy Rasmussen Institute for Genomic Medicine, Abigail Wexner Research Institute, Nationwide Children’s Hospital, Columbus, OH 43205, USA; Department of Pediatrics, The Ohio State University College of Medicine, Columbus, OH 43210, USA

**Keywords:** cranial neural crest cells, *Cse1l*, *Wnt1-Cre*, apoptosis, heart, ventricular myocardium

## Abstract

Human congenital anomalies account for twice the mortality of childhood cancer. Despite advancements in genome sequencing and transgenic mouse models that have aided in understanding their pathogenesis, significant gaps remain. Through a forward genetics approach, we previously discovered the hypo-morphic *anteater* allele of *Cse1l* which displayed variable craniofacial phenotypes. To circumvent the variability seen in this model, we generated a conditional allele of *Cse1l* and genetically ablated it in the dorsal midline giving rise to portions of the nervous system and the cranial neural crest cells using the *Wnt1-Cre*2 driver. Our analysis revealed that *Wnt1-Cre2; Cse1l*^*CRISPR/flox*^ embryos exhibited severe malformations in the forebrain, midbrain, and hindbrain, accompanied by a dramatic hypoplasia of the frontonasal, maxillary, and mandibular processes, and the second pharyngeal arch. *Wnt1-Cre2; Cse1l*^*CRISPR/flox*^ embryos were embryonic lethal by E11.5 likely due to defects in the ventricular myocardium. *Wnt1-Cre2; Cse1l*^*CRISPR/flox*^ embryos exhibited consistently increased apoptosis at E9.5 in the affected tissues along with an increase in p53 expression. These data together show a previously unknown critical function of CSE1L in neural crest cell survival during development.

**Summary Statement:** *Cse1l* is critical for neural crest cell survival and genetic ablation of *Cse1l* in neural crest cells resulted in dramatic apoptosis with increase in p53 expression.

## Introduction

Congenital anomalies affect approximately 5% of the human population and result in twice the annual mortality of childhood cancer (Bower et al., 2010; Stallings et al., 2024). Despite advancements in next-generation sequencing and transgenic models, the genetic, cellular, and molecular mechanisms underlying these malformations remain incompletely understood. In pursuit of characterizing new genes implicated in structural birth defects, we previously used an N-ethyl-N-nitrosourea (ENU) mutagenesis screen and identified *Chromosome Segregation 1 Like (Cse1l)* [previously known as Cellular Apoptosis Susceptibility (*Cas*) and Exportin-2 (*Xpo2*)] to be a regulator of craniofacial development (Blizzard et al., 2021). Based on the appearance of the snout, we named this hypo-morphic allele of *Cse1l as Cse1l*^*anterater*^ (Blizzard et al., 2021). *Cse1l*^*anterater*^ homozygous mutants displayed a remarkably variable phenotypic spectrum, ranging from severe proboscis malformations to near-normal morphology. This spectrum also included anophthalmia or microphthalmia, cleft lip/palate, exencephaly, and polydactyly (Blizzard et al., 2021). These malformations are consistent with the expression of *Cse1l* in the developing forebrain, eyes, tongue, mandibular arch, nasal pits, lateral nasal process and in the future nasal epithelium at E10.5 (Blizzard et al., 2021). However, this variable phenotype precluded any focused molecular investigation of the role for *Cse1l* during development. *Cse1l* has been studied in a number of contexts beyond developmental biology and found to have pleiotropic roles in the cell. It has been implicated in cancer and multiple studies *in vitro* have indicated that loss or reductions in *Cse1l* lead to increased apoptosis and reduced proliferation or cell-cycle arrest (Liao et al., 2008; Lin et al., 2021; Tai et al., 2010; Zhang et al., 2021). Depletion of *Cse1l* is also associated with reduced migration and invasion of cancer cell types (Lorenzato et al., 2013). Interestingly, CSE1L has also been shown to associate with chromatin and regulate the expression of p53 target genes (Tanaka et al., 2007). CSE1L is involved in nuclear import and export of cargo proteins, by recycling importin-α from the nucleus to the cytoplasm for re-use in the next cycle of protein import (Kutay et al., 1997). This activity facilitates import of cargo proteins to maintain repression of several methylated genes. CSE1L is critical for nuclear accumulation of TAZ (Nagashima et al., 2021). TAZ is important for neural crest cell development and nuclear accumulation may change upon loss of CSE1L (Wang et al., 2016; Nagashima et al., 2021). Silencing of *Cse1l* results in failure of nuclear accumulation of HDACs such as HDAC1, HDAC2, and HDAC8, which impact methylation status of the downstream target genes (Dong et al., 2018).

Despite these robust studies of *Cse1l* function in cancer cells *in vitro*, the role of *Cse1l in vivo* and, in particular, during development is largely unknown. This is due to the early embryonic lethality of *Cse1l*^*null/null*^ homozygotes, which die by E5.5 (Bera et al., 2001) Therefore, to understand the role of *Cse1l* in craniofacial development, we generated a novel conditional allele of *Cse1l* and inactivated it in the neural crest cells. Neural crest ablation of *Cse1l* resulted in embryonic lethality at E11.5 and the mutants display hypoplastic facial prominences due to massive apoptosis in the neural crest cells. These data reveal a new *in vivo* function of *Cse1l* during embryonic development.

## Materials and Methods

### Mice and animal husbandry

*Cse1l*^*CRISPR/wt*^ (Blizzard et al., 2021), *Wnt1-Cre* (Danielian et al., 1998), *Wnt1-Cre2* (JAX:022501; Lewis et al., 2013), *Sox2-Cre* (JAX:008454) (Hayashi et al., 2002), *Rosa*^*mTmG*^ (JAX:007676; Muzumdar et al., 2007), *Trp53*^*dell/wt*^ (JAX:002101; Jacks et al., 1994) and a novel *Cse1l*^*flox*^ allele were utilized in this study. *Wnt1-Cre2, Sox2-Cre, Rosa*^*mTmG*^, and *Trp53* mice were obtained from the Jackson laboratory were genotyped by PCR genotyping with specific primer sets as follows: *Cse1l*^*CRISPR/wt*^ F: AGATTCAGAGTCATGGAGCTCA, R: CAATCAGTCAAGGAACAAAGCC; for *Wnt1-Cre2, Wnt1-Cre2* transgene F: CAGCGCCGCAACTATAAGAG, transgene R: CATCGACCGGTAATGCAG, *Wnt1-Cre2* internal positive control F: CAAATGTTGCTTGTCTGGTG, *Wnt1-Cre2* internal positive control R: GTCAGTCGAGTGCACAGTTT; for *Sox2-Cre, Sox2-Cre* F: CTTGTGTAGAGTGATGGCTTGA, *Sox2-Cre* R: TAGTGCCCCATTTTTGAAGG, *Sox2-Cre* transgene: CCAGTGCAGTGAAGCAAATC; for *Rosa*^*mTmG*^, oIMR7318: CTCTGCTGCCTCCTGGCTTCT, oIMR7319: CGAGGCGGATCACAAGCAATA, oIMR7320: TCAATGGGCGGGGGTCGTT; For *Wnt1-Cre*, oIMR1084: GCGGTCTGGCAGTAAAAACTATC, oIMR1085: GTGAAACAGCATTGCTGTCACTT; For *Trp53*, common: TGGATGGTGGTATACTCAGAGC, *Trp53* WT F: AGGCTTAGAGGTGCAAGCTG, and *Trp53* Mut F: CAGCCTCTGTTCCACATACACT. *Trp53*^*delwt*^ mice were maintained in a mixed background of CD1 and C57BL/6J in our laboratory and rest of the breeding was carried out on the C57BL/6J background. Mice were housed with a 12 h light/12 h dark cycle with food and water ad libitum. For timed mating, noon of the day of the vaginal plug was considered as day 0.5. Mouse euthanasia was performed in a carbon dioxide chamber followed by secondary cervical dislocation. All animal work procedures were performed following recommendations in the Guide for Care and Use of Laboratory Animals by the National Institute of Health and approved by the Institutional Animal Care and Use Committee (IACUC) at Nationwide Children’s Hospital (AR21-00067).

### Generation of *Cse1l*^*flox*^ allele

Mice (NCBI Taxon ID 10090) were created at the Cincinnati Children’s transgenic animal and genome editing core (RRID:SCR_022642). The third exon of *Cse1l* was chosen to be surrounded by loxP sites as this will create an early frame shift in the resulting protein. Mouse zygotes (C57BL/6N strain) were injected with 200 ng/μl CAS9 protein (IDT and ThermoFisher), and reagents to insert the 5’ loxP site: 100 ng/μl *Cse1l* 5’ sgRNA (CAGACAACCCTTGTGGAAGG AGG), 75 ng/μl single-stranded donor oligo-nucleotide(GTAGACTTTAGGGACAGTTTAGCTCCTGGGATAGCCTGCTCACAGAAATGAACTCATCAGCACTACAGACAACCCTTGTGGAGTCGACATAACTTCGTATAATGTATGCTATACGAAGTTATAGGAGGCACCTATGCACCTTATCAGTACCATTTAGACATTTCCTGGG AAGC; IDT, Iowa), and the 3’ sgRNA (AGAATTAAAGTAGGTTAAGG TGG) and donor (CAAACTTCATAATTCAGATTGTTAACAGAGTTGAGAGACCTGGGATGTGAAGACAGTCTAAATAAAGAATTAAAGTAGGTTAATAACTTCGTATAATGTATGCTATACGAAGTTATGCTAGCAGGTGGGTATGTGTTGTGTTTATGTCCTCCCAGTTACAACAAGAACC) followed by surgical implantation into pseudo-pregnant female (CD-1 strain) mice. PCR genotyping was performed by amplification of genomic DNA (5’ F: GTCTTAGTTTTGTGGGCTCAGTG, 5’ R: TGAAATGACCTCACTCAGTTCTC, 3’ F: CAGCCATTAATGATGAACAGGTTC, and 3’ R: TGGAATAGGATTTCACTGTGCTTC). A 5’ loxP specific F primer (CTACAGACAACCCTTGTGGAGTCGAC) was used along with the 3’ R primer to test for insertions into the same allele. The PCR products were subject to Sanger Sequencing and pups exhibiting editing of interest were then crossed to wild-type (C57BL/6J) mice and the resulting progeny were Sanger sequenced (CCHMC DNA Sequencing and Genotyping Core) to confirm the alleles generated. Propagation of the *Cse1l*^*flox*^ alleles was done by crossing to wild-type C57BL/6J mice and/or intercross. Further genotyping was with a combination of PCR followed by and/or Sanger sequencing.

### Wnt1-Cre2 and Cse1l^CRISPR^

Given that the *Wnt1Cre2* transgene is integrated into the *E2f1* locus (76.79 cM, 154.4Mb) on chromosome 2 (Dinsmore et al., 2022) which is in the same genomic region as the *Cse1l* gene (87.22 cM, 166.7 Mb) and only 11.5cM (12.3Mb) apart. We crossed the *Cse1l*^*CRISPR/+*^ and *Wnt1-Cre2* alleles to generate a double heterozygous *Cse1l*^*CRISPR/wt*^; *Wnt1-Cre2/wt* animal with the alleles on the same chromosome (cis).

### Western blot

Western blot analysis was performed as previously described (Iyyanar et al., 2022). In brief, craniofacial tissues up to the level of second pharyngeal arch of E9.5 wild-type and *Wnt1-Cre2; Cse1l*^*CRISPR/flox*^ embryos were dissected and snap frozen. Four E9.5 heads were pooled for each genotype as one sample (n=1) and lysed in RIPA buffer containing protease inhibitor cocktail (Thermofisher). Three different biological replicates were assayed for Western blot. Lysates were centrifuged at 21,000 g for 20 min at 4°C and the collected supernatant was quantified using a BCA assay kit (ThermoFisher). 20 µg of protein was separated on a 4–20% mini-PROTEAN TGX stain-free gradient gel (Bio-Rad) and blotted onto a PVDF membrane using Trans-Blot Turbo transfer system. The membranes were blocked with 5% skim milk. Primary antibodies used were anti-CSE1L/CAS rabbit polyclonal (Abcam #151546, 1:1000) and anti-P53 (Cell Signaling 2524S; 1:500). Chemiluminescent detection visualization on a Bio-Rad ChemiDoc with HRP-conjugate secondary antibody anti-rabbit/mouse IgG, (Abcam,1:3000). Proteins were stripped using commercial stripping buffer (Thermofisher). Total protein levels from the stain free gels were used for the normalization of p53 levels. Quantification was carried out using Alpha View software.

### Histology

Histology was performed using Hematoxylin and Eosin as previously described (Blizzard et al., 2021). Embryos were dissected, fixed in 4% paraformaldehyde at 4 C overnight, washed in 70% EtOH, and dehydrated and embedded in paraffin. Blocks were sectioned by microtome at 7 µm, then sections were captured on SuperFrost slides (Cardinal Health, Dublin, OH, USA), and stained with Hematoxylin and Eosin using standard methods. All histological and immunohistochemical studies are performed on at least three animal pairs.

### Whole-mount immunostaining

Wholemount immunostaining was carried out with standard protocols and the embryos were cleared using the iDISCO method (Comai et al., 2020; Renier et al., 2014). Embryos were fixed in 4% PFA overnight at 4°C and serially (25%, 50%, 70%, 100%) passed to 100% methanol and stored at −20°C until further use. The embryos were bleached with 4:1:1 of methanol: hydrogen peroxide: DMSO and then rehydrated serially to PBS. Then the embryos were treated with 0.2% Triton-X-100, followed by permeabilization at 37°C overnight and followed by blocking with 6% goat serum, 10% DMSO, and 0.2% Triton-X-100. Cleaved caspase-3 (Cell signaling, 9661,1:250), and GFP (Avis, GFP-1010,1:500) in 3% goat serum, 5% DMSO, 0.2% Tween-20. After 5 one hour washes, Alexa fluor secondary antibody anti-chicken 488 (Invitrogen, A-11039, 1:500) and anti-rabbit 555 (Invitrogen, A-21428, 1:500) in 3% goat serum was added. After 5 X 1-hour washes, embryos were serially passed to 100% methanol. For iDISCO clearing, embryos were passed to 33% dichloromethane and 66% methanol for 3 hours at room temperature, followed by 100% dichloromethane (Sigma) twice for 15 mins. Finally, the embryos were cleared in dibenzyl ether (100%) (Sigma) twice before imaging in AX-R confocal microscope (Nikon). Z-stack Images were volume rendered and max-IP projected and captured in volume view as snap shots.

### Immunohistochemistry

Embryos were fixed for in 4% PFA at 4°C and passed to 15% and then to 30% sucrose for a day each before the embryo heads were embedded in Optimal Cutting Temperature solution (Sakura) and stored at −80°C. 10 µm sections were obtained on positively charged glass slides with a Leica CM 1860 cryostat. Slides selected for immunohistochemistry were dried, rehydrated with PBS. Antibody retrieval was performed with 0.1 M citrate buffer (pH 6.0) for 15 minutes and cooled down 1.5 hours as previously described (Bittermann et al., 2019). Blocking and secondary antibody processing was as described before (Inskeep et al., 2024). Primary antibody used for p53 (Cell signaling 2524S, 1:200) and secondary antibody anti-mouse Alexa fluor 555 (Invitrogen, A-21422, 1:500). Images were acquired using Zeiss Apotome and p53-positive cells were quantified using ImageJ (red blood cells were subtracted from the p53 staining using non-specific staining images from the far-red channel).

## Results

### Generation and characterization of novel *Cse1l* conditional allele

To investigate the role of *Cse1l* during craniofacial development, we generated a conditional allele of *Cse1l* by inserting two loxP sites flanking exon 3 using CRISPR/CAS9-mediated genome editing (Fig. 1A, B). This is the *Cse1l*^*em2Rstot*^ allele, hereafter referred to as *Cse1l*^*flox*^. We also utilized our previously generated null allele of *Cse1l* (*Cse1l*^*em2Rstot*^, hereafter called *Cse1l*^*CRISPR*^) with a premature stop codon at amino acid 35 (Blizzard et al., 2021). To validate the functionality of these alleles, we crossed the *Wnt1-Cre2; Cse1l*^*CRISPR/wt*^ males to *Cse1l*^*flox/flox*^ female mice and assayed CSE1L protein levels from the cranial region from embryonic day (E)9.5 embryos (Fig. 1C). Western blot analysis using an anti-CSE1L antibody revealed a dramatic reduction in CSE1L protein levels to about 16% of wild-type in *Wnt1-Cre2; Cse1l*^*CRISPR/flox*^ embryos (Fig. 1C). The residual CSE1L protein in mutant embryos is likely attributed to the presence of CSE1L in non-neural crest cells in the cranial region outside the *Wnt1-Cre2* lineage. To further confirm that the *Cse1l* conditional allele is causing the expected disruption to the *Cse1l* locus, we induced a germline deletion of the *Cse1l* conditional allele using the *Sox2-Cre* driver and analyzed the embryos at E8.5 (Fig. 1D, E). Consistent with previously reported *Cse1l-null* embryos, we did not recover any viable *Sox2-Cre2; Cse1l*^*CRISPR/flox*^ mutants in litters dissected at or after E8.5 (n=4 litters, 30 embryos; Chi^2^ test, p=0.023). Prior studies have shown that *Cse1l-null* embryos are typically embryonic lethal around E5.5 (Bera et al., 2001). The only *Sox2-Cre2; Cse1l*^*CRISPR/flox*^ embryo recovered at E8.5 was found to be non-viable (Fig. 1E). These data collectively demonstrate that inactivation of the novel conditional allele of *Cse1l* effectively disrupts CSE1L function.

**Fig. 1.**
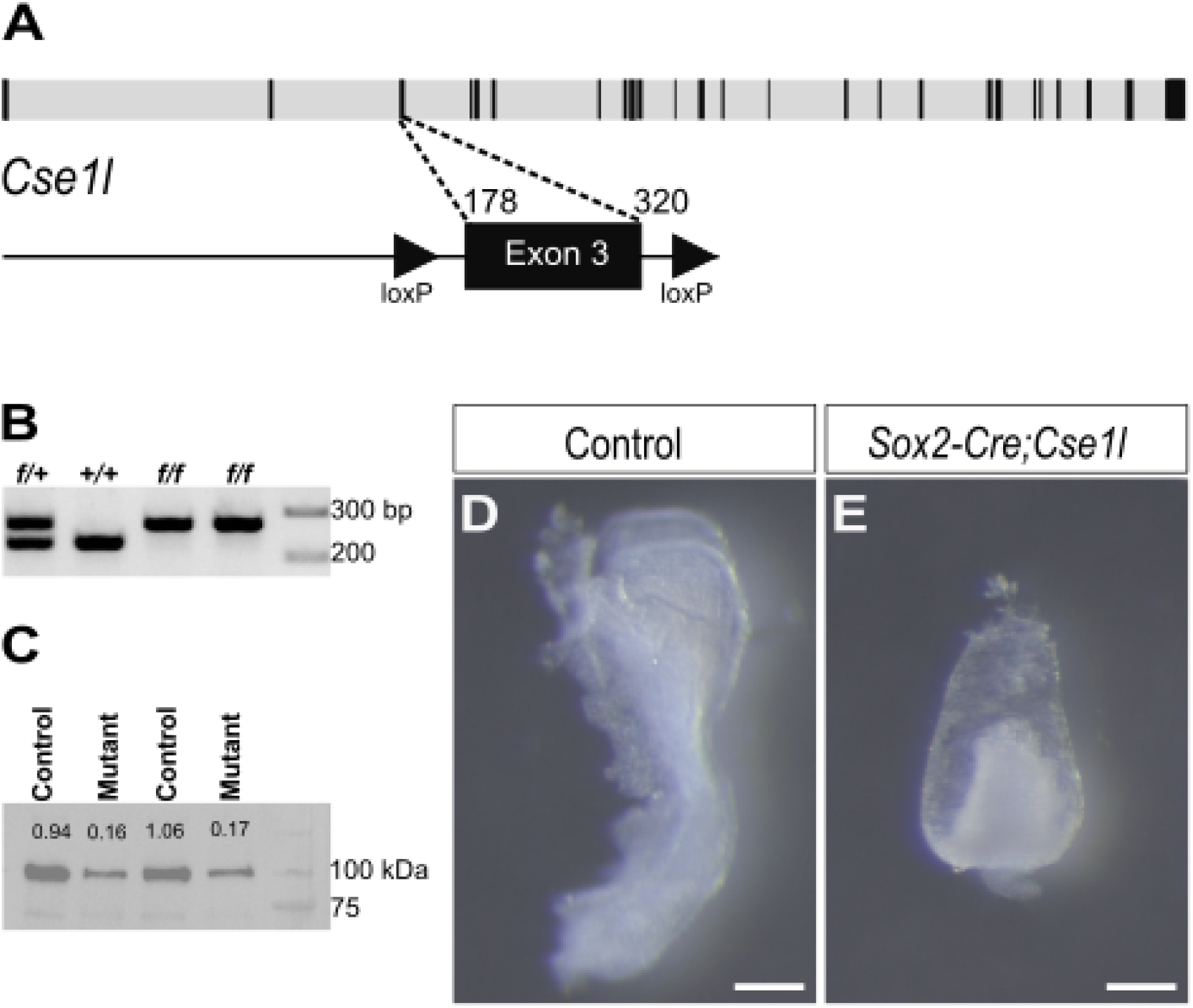
Generation of *Csel1* conditional allele. (A) Schematics of the strategy for generating *Cse1l* conditional allele using a CRISPR/CAS9 genome editing approach. The top row shows the genomic organization of the mouse *Cse1l* locus. Exons 1–25 are indicated with black color boxes. The second row of the schematics shows the insertion of two loxP sites (triangles) flanking exon 3. (B) Representative genotyping gel pictures of *Cse1l*^*flox*^ mice. The amplicons from wildtype and *Cse1l*^*flox*^ alleles are 230 bp and 264 bp, respectively. (C) Western blot analysis using anti-CSE1L antibody indicating a clear reduction of the CSE1L protein in *Wnt1-Cre2; Cse1l*^*CRISPR/flox*^ embryo heads at E9.5 (n=3). (D-E) Lateral view of control (D) and *Sox2-Cre2; Cse1l*^*CRISPR/flox*^ (E) embryos at E8.5.*+/+*, wildtype; *fl/+, Cse1l*^*flox/+*;^ *fl/fl, Cse1l*^*flox/flox*^. Scale bar, 20 µm.

We previously reported that *Cse1l*^*CRISPR/wt*^ heterozygous mice exhibit ocular defects, including microphthalmia, anophthalmia, and cataracts. The most severe phenotype of the *Cse1l*^*anteater/anteater*^ was the proboscis in place of the wild-type snout (Blizzard et al., 2021). During this study, we observed a similar proboscis phenotype in a P0 *Cse1l*^*CRISPR/wt*^ pup, resembling one of the lowly penetrant phenotypes of *Cse1l*^*anteater*^ homozygous mutants (Fig. 2A, B). In litters dissected between E13.5 and E18.5, 50% of *Cse1l*^*CRISPR/wt*^ embryos displayed microphthalmia or an abnormal optic fissure closure defect (15/30; Fig. 2C-F). We also observed mild holoprosencephaly (1/30) in one *Cse1l*^*CRISPR/wt*^ mutant at E13.5 (Fig. 2G, H). In litters dissected between E10.5 and E18.5, we also noted exencephaly (3/129) in *Cse1l*^*CRISPR/wt*^ embryos at E13.5 (Fig. 2I, J). Collectively, these findings demonstrate that haploinsufficiency of *Cse1l* results in a spectrum of craniofacial abnormalities at varying frequencies like the *Cse1l*^*anteater/anteater*^ mutants, highlighting the critical role of CSE1L protein levels in proper craniofacial development.

**Fig. 2.**
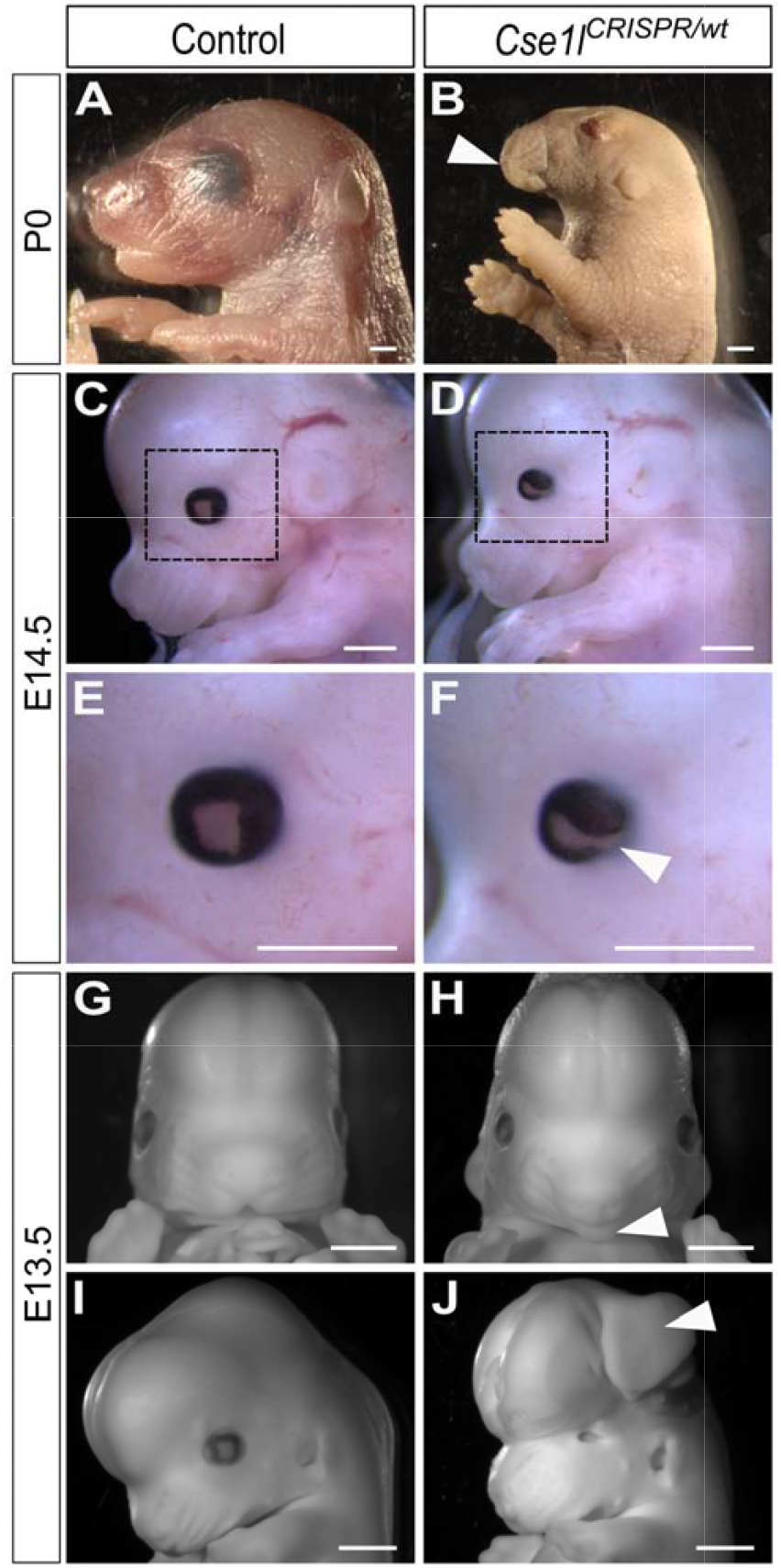
Spectrum of craniofacial phenotype in *Cse1l* heterozygous embryos. Wholemount lateral (A-F, I, J) a**nd** frontal (G, H) views of control (A, C, E, G, I) and *Cse1l*^*CRISPRwt*^ (B, D, F, H, J) embryos at P0 (A, B), E14.5 (C-F), and E13.5 (G-J). White arrowheads in B, F, H, J point to the proboscis-like snout, optic fissure closure defect, holoprosencephaly midface, and exencephaly, respectively, in the *Cse1l* heterozygous mutants. Scale bar, 1 mm.

### Conditional inactivation of *Cse1l* in neural crest cells (NCCs) results in embryonic lethality at E11.5 and truncated craniofacial tissues

To investigate the role of *Cse1l* in neural crest cells (NCCs), we inactivated *Cse1l* using the *Wnt1-Cre2* driver (Lewis et al., 2013). At E11.5, *Wnt1-Cre2; Cse1l*^*CRISPR/flox*^ embryos exhibited severe malformations in the forebrain, midbrain, and hindbrain regions, accompanied by a dramatic hypoplasia of the frontonasal, maxillary, and mandibular processes, as well as the second pharyngeal arch (Fig. 3A, B). Notably, *Wnt1-Cre2; Cse1l*^*CRISPR/flox*^ embryos lacked a heartbeat and were dead by E11.5. During this study, emerging reports indicated that the *Wnt1-Cre2* transgene is inserted into the *E2f1* locus in chromosome 2 and causes ectopic Cre activation in spermatogonia of adult male mice and in the definitive endoderm (Dinsmore et al., 2022). To further verify that the abnormalities observed in the *Wnt1-Cre2; Cse1l*^*CRISPR/flox*^ embryos are due to the loss of *Cse1l* in specifically in neural crest cells, we conditionally inactivated *Cse1l* using the *Wnt1-Cre* driver (*Wnt-1/GAL4/Cre-11*; Jax:003829) (Danielian et al., 1998). As expected, *Wnt1-Cre; Cse1l*^*CRISPR/flox*^ embryos also exhibited embryoni**c** lethality at E11.5, lacking a heartbeat. These embryos also displayed severe hypoplasia of the frontonasal, maxillary, mandibular, and second pharyngeal arches (Fig. 3C, D). Given the striking phenotypic similarity between the two *Wnt1-Cre* driver lines and the consistent embryonic lethality due to cardiac malformations, we conclude that the phenotype of *Wnt1-Cre2; Cse1l*^*CRISPR/flox*^ is likely due to the loss of *Cse1l* in the neural crest cells and any potential phenotypes due effects of the Cre alleles are unlikely to significantly impact our understanding of the role of *Cse1l* in neural crest development. Therefore, we proceeded with further analyses using the *Wnt1-Cre2; Cse1l*^*CRISPR/flox*^ mutants. Collectively, these data demonstrate that *Cse1l* function within NCCs is essential for proper heart and craniofacial development.

**Fig. 3.**
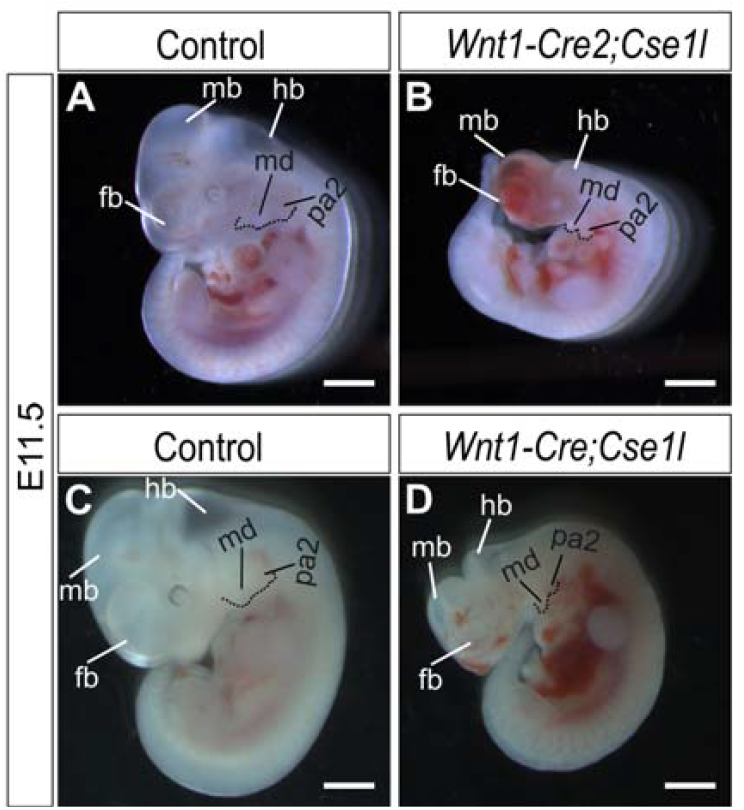
Craniofacial defects and embryonic lethality in the *Cse1l* neural crest mutants. (A-D) Lateral view of control (A, C) and *Wnt1-Cre2; Cse1l*^*CRISPR/flox*^ (B) and *Wnt1-Cre; Cse1l*^*CRISPR/flox*^ (D) embryos at E11.5. fb, forebrain; hb, hindbrain; mb, midbrain; md, mandibular arch; pa2, second pharyngeal arch. Scale bar, 1 mm.

### Cardiac malformations in the *Cse1l* neural crest mutants

Stottmann et al., previously demonstrated that the cardiac neural crest cells are critical for the proper maturation and thickening of the ventricular myocardium (Stottmann et al., 2004). To further explore the cardiac phenotype of the *Cse1l* neural crest mutants, we performed a histological analysis of the heart in the transverse plane at E10.5 and E11.5. At E10.5, the ventricular myocardium looked comparable between the control and *Wnt1-Cre2; Cse1l*^*CRISPR/flox*^ mutants (Fig. 4A-D). However, by E11.5, while control embryos showed progressive ventricular myocardium thickening, the *Wnt1-Cre2; Cse1l*^*CRISPR/flox*^ mutants exhibite**d** a marked reduction in condensed myocardium (Fig. 4E-H). Notably, we observed consistent hemorrhage within the ventricular chambers of developing hearts of *Wnt1-Cre2;Cse1l*^*CRISPR/flox*^ mutants (Fig. 4E, F). These data are consistent with the previously documented phenotype of neural crest mutants of *Bmpr1a*, where a lack of neural crest-derived signals from the epicardium is thought to result in failure of the ventricular myocardium (Stottmann et al., 2004).

**Fig. 4.**
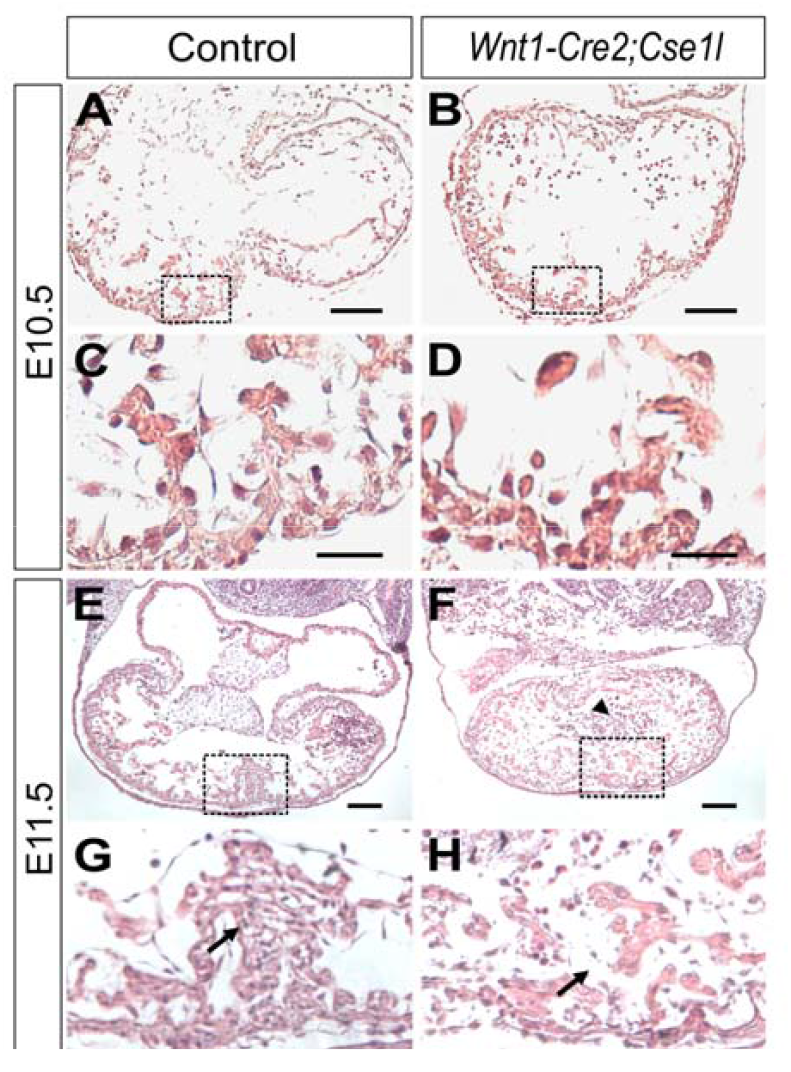
Cardiac defects in *Wnt1-Cre2; Cse1l*^*CRISPR/flox*^ embryos. (A-H) Hematoxylin and Eosin staining in the transverse sections of control (A, C, E, G) and *Wnt1-Cre2*; *Cse1l*^*CRISPR/flox*^ (B, D, F, H) embryos at E10.5 (A-D), and E11.5 (E-H). Arrows in G, H point to the interventricular septum and arrowhead in F point to the hemorrhage in the ventricles of *Wnt1-Cre2;Cse1l*^*CRISPR/flox*^ embryos. n=3 each at E10.5 and E11.5. Scale bars in C, D, G, H are 25 µm and in A, B, E, F are 100 µm.

### *Wnt1-Cre2; Cse1l*^*CRISPR/flox*^ embryos lack neural crest cells migrating into the outflow tract and epicardium

To investigate whether a lack of neural crest migration or apoptosis is responsible for the cardiac phenotype observed in the *Wnt1-Cre2; Cse1l*^*CRISPR/flox*^ mutants, we carried out neural crest cell lineage tracing using the *Wnt1-Cre2* allele in concert with the *Rosa*^*mTmG*^ reporter (Muzumdar et al., 2007). We followed this with wholemount immunostaining for GFP and Cleaved Caspase-3 in control and *Wnt1-Cre2; Cse1l*^*CRISPR/flox*^; *Rosa*^*mTmG/wt*^ mutants at E10.5. In the control embryos, previously observed neural crest lineages were noticed in the outflow tract and ventricular region (Fig. 5A, C, E) (Kirby et al., 1983). In contrast, the *Wnt1-Cre2; Cse1l*^*CRISPR/flox*^; *Rosa*^*mTmG/wt*^ mutants exhibited a complete lack of reporter activity in both outflow tract and ventricular region in lateral, transverse and frontal planes (Fig. 5B, D, F). However, we did not notice any Cleaved Caspase-3 labeling in the developing heart of control or *Wnt1-Cre2; Cse1l*^*CRISPR/flox*^ mutants at E10.5 (Fig. 5A-F). These data together show that the lack of neural crest cells may have contributed to the lack of thickening of the ventricular myocardium in *Wnt1-Cre2; Cse1l*^*CRISPR/flox*^ mutants at E11.5 as has been seen before in other neural crest conditional mutants (Stottmann et al., 2004).

**Fig. 5.**
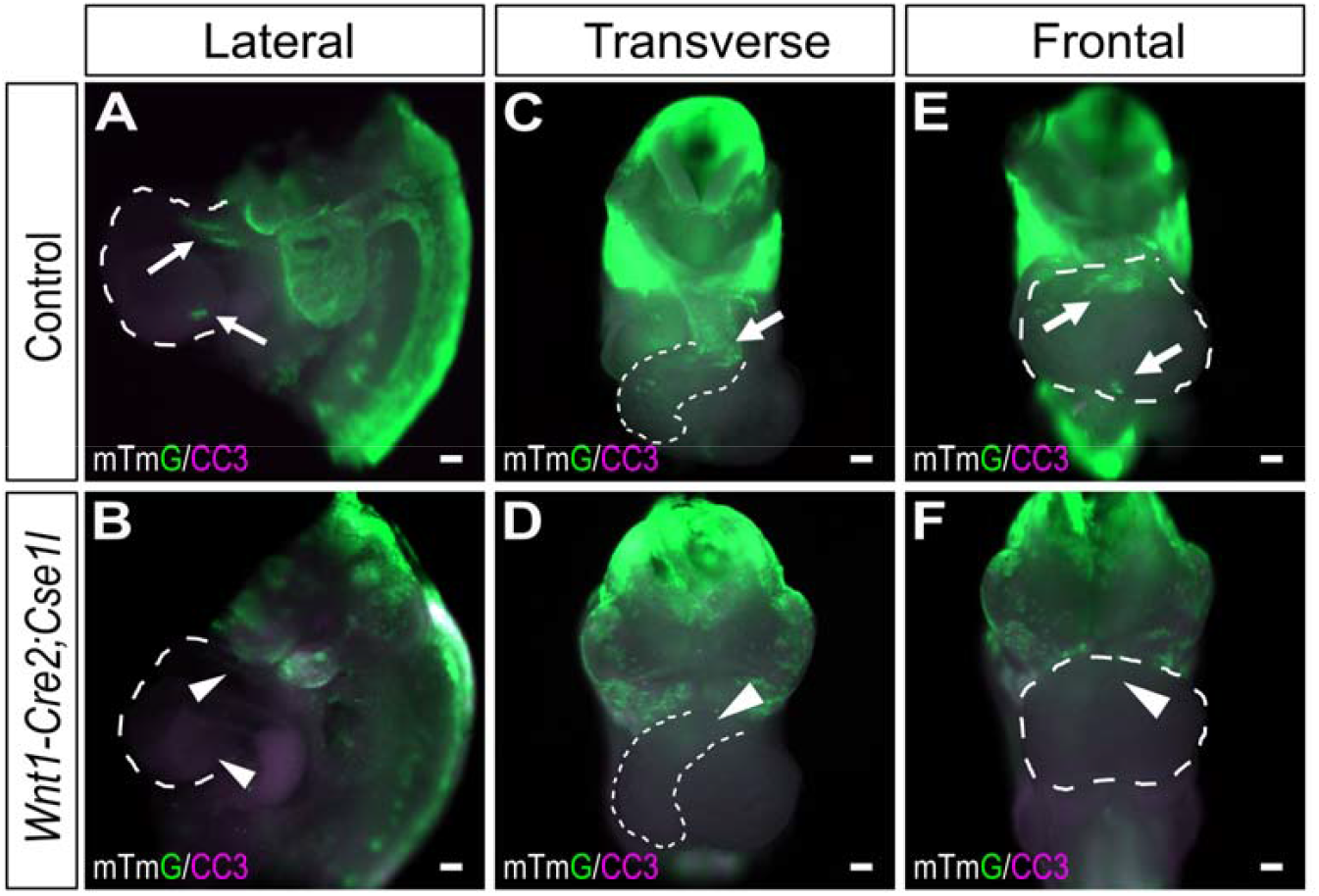
Lack of cardiac neural crest cell in the outflow tract and ventricle in *Wnt1-Cre2; Cse1l*^*CRISPR/flox*^ embryos. (A-F) Wholemount immunostaining for *Wnt1-Cre2; Rosa*^*mTmG*^ (GFP) (green) and Cleaved Caspase-3 (CC3, purple) in control (A, C, E) and *Wnt1-Cre2; Cse1l*^*CRISPR/flox*^ (B, D, F) hearts in lateral (A, B), transverse (C, D), and frontal planes (E, F). Arrows point to the neural crest cells migrated into the outflow tract and ventricles in control embryos and arrowheads point to the lack of neural crest cells in the outflow tract and ventricles in *Wnt1-Cre2; Cse1l*^*CRISPR/flox*^ embryos. n=3. Scale bar, 100 µm.

### *Cse1l* is required for the survival of cranial neural crest cells during development

To determine whether the severe craniofacial phenotype observed in *Wnt1-Cre2; Cse1l*^*CRISPR/flox*^ embryos resulted from cell death, we performed whole-mount immunostaining for cleaved Caspase 3 in E9.5 embryos. Embryos were cleared using the iDISCO clearing method and imaged with confocal microscopy. While the control embryos exhibited very little apoptosis in the embryonic head (Fig. 6A), extensive apoptosis in the mid-hindbrain region and in the craniofacial region was observed in the *Wnt1-Cre2; Cse1l*^*CRISPR/flox*^ mutants along the proximal-distal and lateral-medial axes at E9.5 (Fig. 6B-F). These findings demonstrate that *Cse1l* is critical for the neural crest cell survival during development and inactivation of *Cse1l* in the neural crest cells leads to increase in programmed cell death ultimately resulting in severe craniofacial defects.

**Fig. 6.**
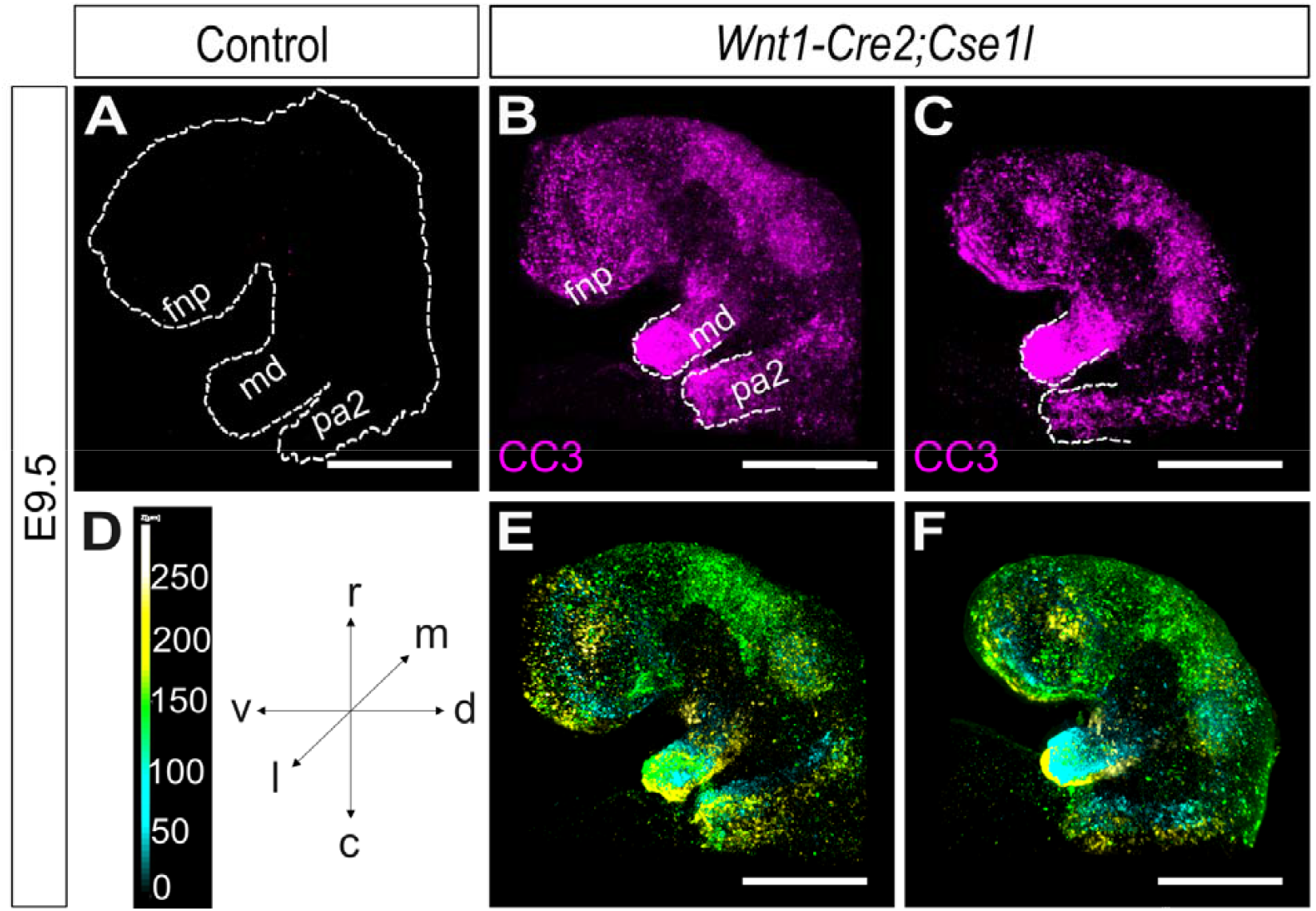
Massive apoptosis in the craniofacial region of *Wnt1-Cre2; Cse1l*^*CRISPR/flox*^ embryos. (A-C) Wholemount view of three dimensional confocal imaging for Cleaved Caspase-3 (CC3, purple) in control (A) and *Wnt1-Cre2; Cse1l*^*CRISPR/flox*^ (B, C) embryos at E9.5. (D-F) Maximum intensity projections and depth coding of CC3 immunostaining in *Wnt1-Cre2; Cse1l*^*CRISPR/flox*^ embryos. Values in D represent the distance in the z plane in µm. n=4. c, caudal; d, dorsal; fnp, frontonasal process; l, lateral; m, medial; md, mandibular process; pa2, second pharyngeal arch; r, rostral; v, ventral. Scale bar, 100 µm.

### Increased P53 expression in the *Wnt1-Cre2; Cse1l*^*CRISPR/flox*^ embryos

placode, mandible, and second pharyngeal arch regions of *Wnt1-Cre2; Cse1l*^*CRISPR/flox*^ embryos compared to controls (Fig. 7A, B) Quantification revealed a significant increase in the number of P53-positive cells within the craniofacial region of *Wnt1-Cre2; Cse1l*^*CRISPR/flox*^ embryos (p < 0.0001, Fig. 7C). Western blot analysis using an anti-P53 antibody revealed an CSE1L was shown to bind to P53 and inhibit P53 target genes by affecting their methylation status (Tanaka et al., 2007). To investigate the potential involvement of P53 signaling in the observed programmed cell death, we performed immunofluorescence staining for P53 in sagittal sections of control and *Wnt1-Cre2; Cse1l*^*CRISPR/flox*^ embryo heads at E9.5. Increased P53 immunoreactivity was observed in the optic 80% increase in the expression of P53 in embryonic head samples from *Wnt1-Cre2; Cse1l*^*CRISPR/flox*^ embryos at E9.5 (Fig. 7D). These data together suggest that apoptosis noticed in the *Wnt1-Cre2;Cse1l*^*CRISPR/flox*^ embryos could be attributed to aberrant P53 signaling.

**Fig. 7.**
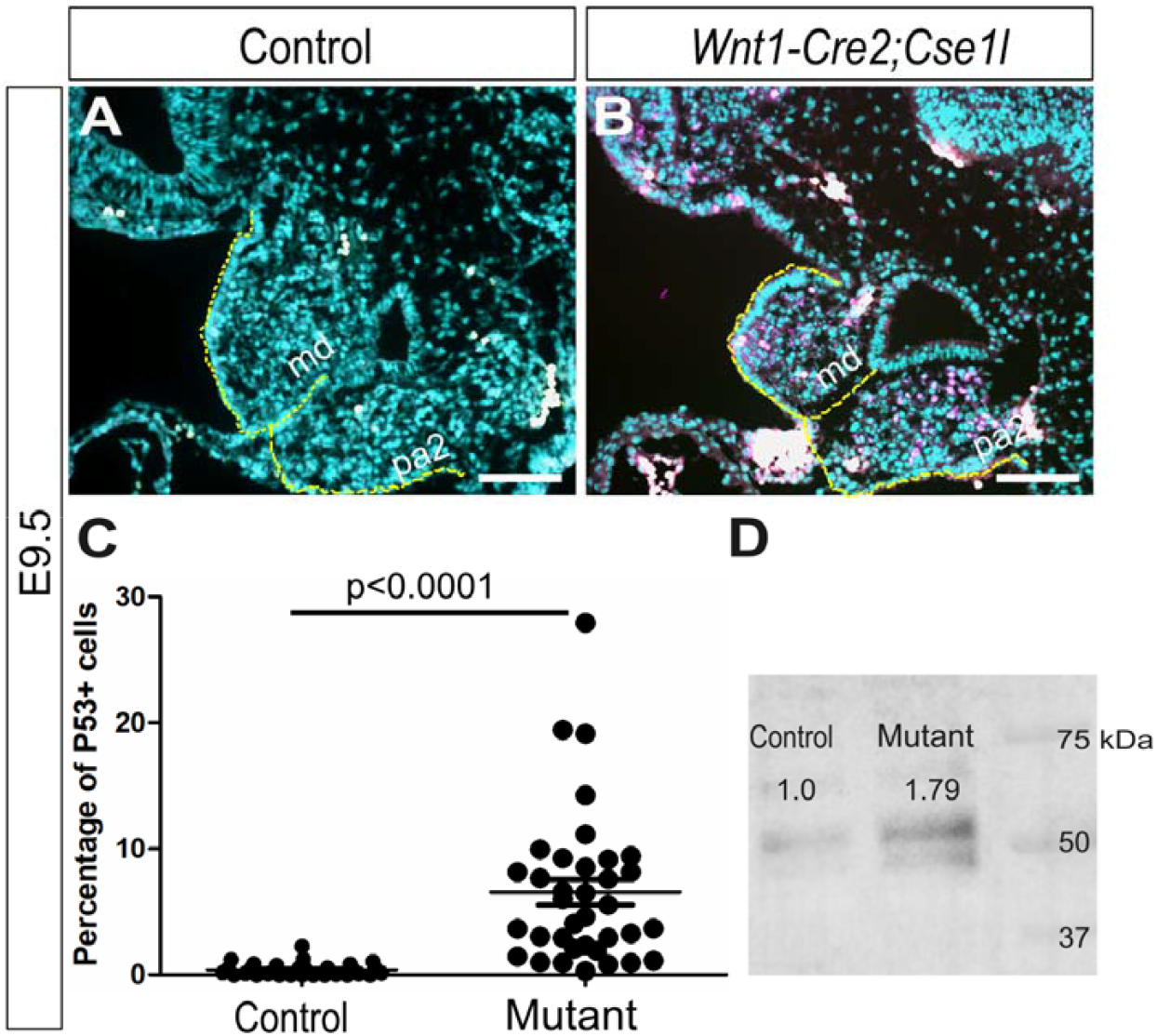
Increased P53 expression in *Wnt1-Cre2; Cse1l*^*CRISPR/flox*^ embryos. (A-B) Immunostaining for P53 in sagittal sections of control (A) and *Wnt1-Cre2; Cse1l*^*CRISPR/flox*^ (B) embryo heads at E9.5. P53 positive cells and red blood cells are labelled in violet and white, respectively. Quantification of P53+ cells (C) in control and *Wnt1-Cre2; Cse1l*^*CRISPR/flox*^ embryo heads and unpaired t-test was performed for statistical analysis. Western blot analysis (D) against anti-p53 antibody in control and *Wnt1-Cre2; Cse1l*^*CRISPR/flox*^ heads at E9.5. N=3. md, mandibular arch, pa2, second pharyngeal arch. Scale bar, 100 µm.

Previous reports have shown that haploinsufficiency of *Trp53* rescued the severe apoptosis and craniofacial phenotype of *Tcof1*^*null/wt*^ and *Twsg1* ^*null/wt*^ mutants (Billington et al., 2011; Jones et al., 2008). To explore the potential role of p53-mediated apoptosis in the craniofacial defects observed in *Wnt1-Cre2; Cse1l*^*CRISPR/flox*^, we attempted to rescue the phenotype by genetically reducing *Trp53* dosage. We crossed *Cse1l*^*flox/wt*^; *Trp53* ^*del/wt*^ or *Cse1l*^*flox/flox*^; *Trp53*^*del/wt-*^ mice with *Cse1l*^*CRISPR/wt*^; *Wnt1-Cre2* mice and analyzed the phenotype of the resulting embryos at E10.5 (Fig. 8). Of the 19 *Wnt1-Cre2; Cse1l*^*CRISPR/flox*^; *Trp53* ^*del/wt*^ embryos examined, one exhibited a complete rescue of the craniofacial defects, displaying a phenotype comparable to wild-type controls, while 18 out of 19 mutants resembled the phenotype of *Wnt1-Cre2; Cse1l*^*CRISPR/flox*^ embryos (Fig. 8C, D). Next, we investigated whether complete loss of *Trp53* results in further rescue of *Wnt1-Cre2; Cse1l*^*CRISPR/flox*^ embryos b**y** crossing *Wnt1-Cre2*; *Csel1*^*CRISPR/wt*^; *Trp53*^*del/+*^ females to *Cse1l*^*flox/flox*^; *Trp53*^*del/del*^ males and analyzed the resulting embryos at E10.5. We observed three out of eight *Cse1l*^*flox/wt*^; *Trp53*^*del/del*^ embryos exhibited exencephaly along the fore, mid, and hindbrain regions, a phenotype that was previously observed in the *Trp53*^*del/del*^ embryos (Figure 8F, G) (Rinon et al., 2011). Interestingly, *Wnt1-Cre2; Cse1l*^*CRISPR/flox*^; *Trp53*^*del/del*^ embryos did not exhibit rescue in the craniofacial phenotypes caused due to loss of *Cse1l* function in the neural crest cells (Figure 8H). These data together indicate that *Cse1l* might function through other signaling pathways in addition to p53 to regulate cell survival, however the role of *Trp53* in the *Cse1l* phenotype cannot be ruled out

**Fig. 8.**
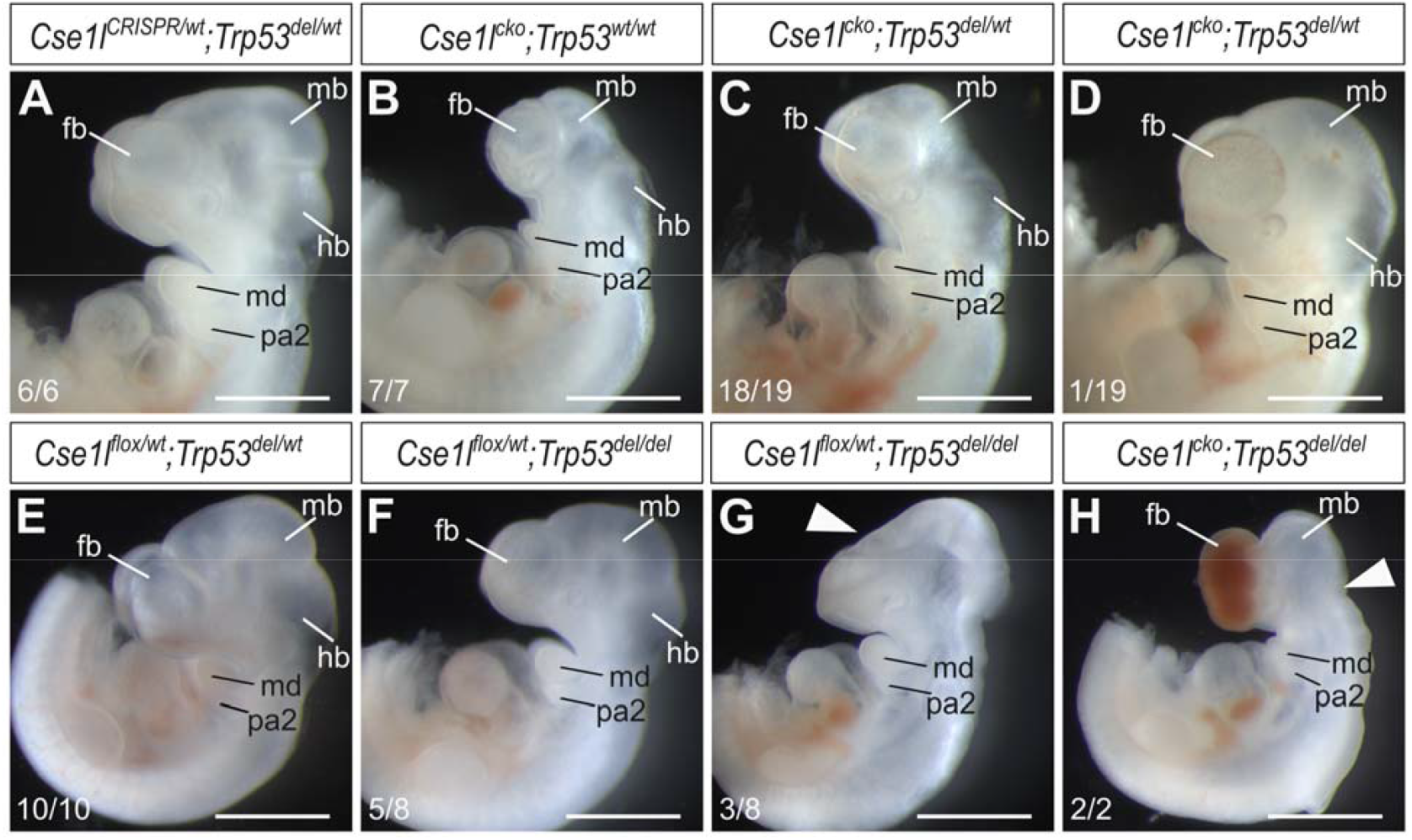
Loss of *Trp53* fails to rescue the phenotype in most of *Wnt1-Cre2; Cse1l*^*CRISPR/flox*^ embryos. (A-D) Wholemount lateral views of *Wnt1-Cre2; Cse1l*^*CRISPR/wt*^*;Trp53*^*del/wt*^ (A), *Wnt1-Cre2; Cse1l*^*CRISPR/flox*^ *Trp53*^*wt/wt*^ (B), *Wnt1-Cre2; Cse1l*^*CRISPR/flox*^*;Trp53*^*del/wt*^ (C, D), *Cse1l*^*flox/wt*^*;Trp53*^*del/wt*^ (E), *Cse1l*^*flox/wt*^*;Trp53*^*del/del*^ (F, G), and *Wnt1-Cre2; Cse1l*^*CRISPR/flox*^*;Trp53*^*del/del*^ (H) at E10.5. Arrowhead in G points to the exencephaly in the entire developing neurocranium of *Cse1l*^*flox/wt*^; *Trp53*^*del/del*^ embryos and arrowhead in H points to the exencephaly in the hindbrain region of *Wnt1-Cre2; Cse1l*^*CRISPR/flox*^*;Trp53*^*del/del*^ embryos. fb, forebrain; hb, hindbrain; mb, midbrain; md, mandibular arch; pa2, second pharyngeal arch. Scale bar, 1 mm.

## Discussion

The essential role of *Cse1l* during embryonic development has been challenging to elucidate due to the early embryonic lethality of *Cse1l*^*null/null*^ mutants at E5.5 (Bera et al., 2001). Our previous work identified an ENU-induced hypo-morphic allele, *Cse1l*^*anteater*^, which exhibited highly variable phenotypes, limiting in-depth molecular and cellular analysis (Blizzard et al., 2021). To overcome these limitations, we report here a novel conditional allele of *Cse1l*. Utilizing this tool, we investigated the specific function of CSE1L within the neural crest-derived craniofacial region. Conditional ablation of *Cse1l* in the neural crest mesenchyme resulted in severe developmental defects, including the loss of midbrain, hindbrain, and facial processes such as the mandibular arch, frontonasal process, and maxillary process due to a massive increase in apoptosis by E9.5. Collectively, these findings reveal a previously unrecognized critical role for *Cse1l* in the survival of cranial neural crest cells. *Wnt1-Cre2; Cse1l*^*CRISPR/flox*^ embryos displayed a substantial deficit in neural crest cells to both the outflow tract and the ventricular region by E10.5. This lack of neural crest cells resulted in the lack of ventricular myocardium thickening by E11.5, which we suggest ultimately leads to embryonic lethality. Notably, Stottmann et al., reported a remarkably similar phenotype in *Wnt1-Cre; Bmpr1a*^*flox/null*^ embryos (Stottmann e**t** al., 2004). More recently, Tang et al. utilized *Wnt1-Cre* lineage tracing in mice and replication-incompetent retrovirus labeling in chick embryos to demonstrate that cardiac neural crest cells contribute to the epicardium and the underlying ventricular myocardium (Tang et al., 2019). This contribution of cardiac neural crest cells to the ventricular myocardium is intriguing. Future investigations employing single-cell transcriptomics combined with refined lineage tracing techniques will be crucial in determining whether neural crest cells in the epicardium non-cell-autonomously signal to the myocardium for proper heart morphogenesis, or if cardiac neural crest cells directly integrate into and form a portion of the ventricular myocardium.

CSE1L plays a crucial role in nuclear transport, mediating various cellular programs and signal transduction pathways. It may govern neural crest development by facilitating the nuclear import and export of specific cargo molecules. Specifically, CSE1L participates in the IMPORTIN-α transport cycle by binding to IMPORTIN-α in the nucleus after cargo release. This binding displaces IMPORTIN-β, enabling IMPORTIN-α export to the cytoplasm (Kutay et al., 1997). This CSE1L-mediated IMPORTIN-α shuttling is essential for the nuclear accumulation of TAZ (Nagashima et al., 2021). Interestingly, neural crest cell ablation of *Yap* and *Taz* (*Wnt1-Cre; Yap*^*flox/flox*^; *Taz*^*flox/flox*^) resulted in aberrant cell death in the mandibular arch and in the optic placode at E9.5 and these *Wnt1-Cre; Yap*^*flox/flox*^; *Taz*^*flox/flox*^ embryos are dead by E10.5 due to cardiac malformations (Wang et al., 2016). Given the striking similarity between the *Cse1l* and *Yap/Taz* mutants, the craniofacial defects observed in our *Cse1l* neural crest mutants could be attributed to disrupted TAZ localization. Furthermore, siRNA-mediated silencing of *Cse1l* in B2-1 cells altered the expression of various transcription factors and activated silenced genes, not through methylation changes, but by delocalizing HDAC1, 2, and 8 to the cytosol (Dong et al., 2018). Therefore, loss of CSE1L may lead to aberrant gene expression patterns in developing neural crest cells, ultimately compromising cell survival. Future studies will be directed towards elucidating specific targets of CSE1L in nuclear transport during embryonic development.

CSE1L has been implicated in the P53 pathway, where it appears to inhibit the expression of *Trp53* target genes through direct binding with P53. Notably, *Cse1l* silencing leads to increased H3K27 methylation at the *Pig3* locus, a *Trp53* target gene (Tanaka et al, 2007). This observation aligns with our findings in *Wnt1-Cre2; Cse1l*^*CRISPR/flox*^ mutants, where loss of *Cse1l* resulted in elevated apoptosis. It remains to be identified whether CSE1L directly binds to chromatin regions with p53 to regulate target gene expression. While several studies have confirmed CSE1L’s role in modulating target gene methylation indirectly through HDAC translocation into the nucleus, the direct DNA binding function for inhibitory gene expression remains less explored (Dong et al., 2018).

Interestingly, IMPORTIN-α3 binding to the P53 protein isn critical for the nuclear localization under stress-induced conditions (Marchenko et al., 2010). We observed strong P53 expression in the mandibular and second pharyngeal arches, as well as the frontonasal region in our *Wnt1-Cre2; Cse1l*^*CRISPR/flox*^ mutants at E9.5. To investigate whether aberrant P53 signaling contributes to the increased cell death like in the *Tcof*^*wt/null*^ and *Twsg1*^*null/null*^ mutants, we attempted to rescue the phenotype by removing one allele of *Trp53* in *Wnt1-Cre2; Cse1l*^*CRISPR/flox*^ mutants (Billington et al., 2011; Jones et al., 2008). While most *Wnt1-Cre2; Cse1l*^*CRISPR/flox*^*;Trp53*^*del/wt*^ embryos (18/19) still exhibited craniofacial deficits similar to *Wnt1-Cre2;Cse1l*^*CRISPR/flox*^ mutants, one embryo showed a phenotypic rescue, resembling a wild-type control. However, *Wnt1-Cre2*; *Cse1l*^*CRISPR/flox*^; *Trp53*^*del/del*^ embryos did not exhibit any rescue in the phenotype of *Wnt1-Cre2*; *Cse1l*^*CRISPR/flox*^ embryos. Consequently, while P53 upregulation could be a secondary effect of *Cse1l* loss, a functional role for the P53 pathway downstream of *Cse1l* cannot be excluded. The variability in the *Cse1*^*anteater*^ and *Cse1l*^*CRISPR/wt*^ phenotypes as well as a massive neural crest cell death and early embryonic lethality in the *Wnt1-Cre2; Cse1l*^*CRISPR/flox*^ mutants have hindered a complete mechanistic understanding (Blizzard et al., 2021). Indeed, RNA-seq analysis in *Cse1*^*anteater*^ mutant heads did not reveal any differentially regulated genes. We suspect this was due to the wide phenotypic spectrum in mutants, which did not allow any gene or pathway to rise above the background (Blizzard et al., 2021). However, future studies focusing on CSE1L’s role in nuclear transport regulation may significantly advance our knowledge of its molecular function.

## Acknowledgements

We appreciate comments on the manuscript from the Stottmann laboratory group. We thank Dr. Samantha A. Brugmann, Cincinnati Children’s Hospital Medical Center for sharing the *Wnt1-Cre* mice (*Wnt-1/GAL4/Cre-11)*. We thank Nicole Constantino for help with maintaining the *Cse1l* mouse line in our laboratory. We thank the Cincinnati Children’s Transgenic and Genomic Editing Core and Yueh-Chiang Hu for their support in creating the *Cse1l*^*flox*^ line. We thank Dr. Tatyana Vetter, Director of the Microscopy core, Nationwide Children’s Hospital for help with confocal imaging. Funding for this project comes from the National Institutes of Health, R01DE027091, and Abigail Wexner Research Institute at Nationwide Children’s Hospital recruitment funds (R.W.S.). No competing interests are declared.

## Reference List

Bera, T. K., Bera, J., Brinkmann, U., Tessarollo, L., & Pastan, I. (2001). Cse1l is essential for early embryonic growth and development. Mol Cell Biol, 21(20), 7020–7024. 10.1128/MCB.21.20.7020-7024.2001

Billington, C. J., Jr., Ng, B., Forsman, C., Schmidt, B., Bagchi, A., Symer, D. E.,… Petryk, A. (2011). The molecular and cellular basis of variable craniofacial phenotypes and their genetic rescue in Twisted gastrulation mutant mice. Dev Biol, 355(1), 21–31. 10.1016/j.ydbio.2011.04.026

Bittermann, E., Abdelhamed, Z., Liegel, R. P., Menke, C., Timms, A., Beier, D. R., & Stottmann, R. W. (2019). Differential requirements of tubulin genes in mammalian forebrain development. PLoS Genet, 15(8), e1008243. 10.1371/journal.pgen.1008243

Blizzard, L. E., Menke, C., Patel, S. D., Waclaw, R. R., Lachke, S. A., & Stottmann, R. W. (2021). A Novel Mutation in Cse1l Disrupts Brain and Eye Development with Specific Effects on Pax6 Expression. J Dev Biol, 9(3). 10.3390/jdb9030027

Bower, C., Rudy, E., Callaghan, A., Quick, J., & Nassar, N. (2010). Age at diagnosis of birth defects. Birth Defects Res A Clin Mol Teratol, 88(4), 251–255. 10.1002/bdra.20658

Comai, G. E., Tesarova, M., Dupe, V., Rhinn, M., Vallecillo-Garcia, P., da Silva, F.,… Tajbakhsh, S. (2020). Local retinoic acid signaling directs emergence of the extraocular muscle functional unit. PLoS Biol, 18(11), e3000902. 10.1371/journal.pbio.3000902

Danielian, P. S., Muccino, D., Rowitch, D. H., Michael, S. K., & McMahon, A. P. (1998). Modification of gene activity in mouse embryos in utero by a tamoxifen-inducible form of Cre recombinase. Current Biology, 8(24), 1323–S1322. 10.1016/S0960-9822(07)00562-3

Dinsmore, C. J., Ke, C. Y., & Soriano, P. (2022). The Wnt1-Cre2 transgene is active in the male germline. Genesis, 60(3), e23468. 10.1002/dvg.23468

Dong, Q., Li, X., Wang, C. Z., Xu, S., Yuan, G., Shao, W.,… Zhu, B. (2018). Roles of the CSE1L-mediated nuclear import pathway in epigenetic silencing. Proc Natl Acad Sci U S A, 115(17), E4013–E4022. 10.1073/pnas.1800505115

Hayashi, S., Lewis, P., Pevny, L., & McMahon, A. P. (2002). Efficient gene modulation in mouse epiblast using a Sox2Cre transgenic mouse strain. Mechanisms of Development, 119, S97–S101. 10.1016/S0925-4773(03)00099-6

Inskeep, K. A., Crase, B., Dayarathna, T., & Stottmann, R. W. (2024). SMPD4-mediated sphingolipid metabolism regulates brain and primary cilia development. Development, 151(22). 10.1242/dev.202645

Iyyanar, P. P. R., Wu, Z., Lan, Y., Hu, Y. C., & Jiang, R. (2022). Alx1 Deficient Mice Recapitulate Craniofacial Phenotype and Reveal Developmental Basis of ALX1-Related Frontonasal Dysplasia. Front Cell Dev Biol, 10, 777887. 10.3389/fcell.2022.777887

Jacks, T., Remington, L., Williams, B. O., Schmitt, E. M., Halachmi, S., Bronson, R. T., & Weinberg, R. A. (1994). Tumor spectrum analysis in p53-mutant mice. Current Biology, 4(1), 1–7. 10.1016/S0960-9822(00)00002-6

Jones, N. C., Lynn, M. L., Gaudenz, K., Sakai, D., Aoto, K., Rey, J. P.,… Trainor, P. A. (2008). Prevention of the neurocristopathy Treacher Collins syndrome through inhibition of p53 function. Nat Med, 14(2), 125–133. 10.1038/nm1725

Kutay, U., Bischoff, F. R., Kostka, S., Kraft, R., & Görlich, D. (1997). Export of Importin α from the Nucleus Is Mediated by a Specific Nuclear Transport Factor. Cell, 90(6), 1061–1071. 10.1016/S0092-8674(00)80372-4

Lewis, A. E., Vasudevan, H. N., O’Neill, A. K., Soriano, P., & Bush, J. O. (2013). The widely used Wnt1-Cre transgene causes developmental phenotypes by ectopic activation of Wnt signaling. Dev Biol, 379(2), 229–234. 10.1016/j.ydbio.2013.04.026

Liao, C. F., Luo, S. F., Li, L. T., Lin, C. Y., Chen, Y. C., & Jiang, M. C. (2008). CSE1L/CAS, the cellular apoptosis susceptibility protein, enhances invasion and metastasis but not proliferation of cancer cells. J Exp Clin Cancer Res, 27(1), 15. 10.1186/1756-9966-27-15

Lin, H. C., Li, J., Cheng, D. D., Zhang, X., Yu, T., Zhao, F. Y.,… Yao, M. (2021). Nuclear export protein CSE1L interacts with P65 and promotes NSCLC growth via NF-kappaB/MAPK pathway. Mol Ther Oncolytics, 21, 23–36. 10.1016/j.omto.2021.02.015

Lorenzato, A., Biolatti, M., Delogu, G., Capobianco, G., Farace, C., Dessole, S.,… Di Renzo, M. F. (2013). AKT activation drives the nuclear localization of CSE1L and a pro-oncogenic transcriptional activation in ovarian cancer cells. Exp Cell Res, 319(17), 2627–2636. 10.1016/j.yexcr.2013.07.030

Marchenko, N. D., Hanel, W., Li, D., Becker, K., Reich, N., & Moll, U. M. (2010). Stress-mediated nuclear stabilization of p53 is regulated by ubiquitination and importin-alpha3 binding. Cell Death Differ, 17(2), 255–267. 10.1038/cdd.2009.173

Muzumdar, M. D., Tasic, B., Miyamichi, K., Li, L., & Luo, L. (2007). A global double-fluorescent Cre reporter mouse. Genesis, 45(9), 593–605. 10.1002/dvg.20335

Nagashima, S., Maruyama, J., Honda, K., Kondoh, Y., Osada, H., Nawa, M.,… Hata, Y. (2021). CSE1L promotes nuclear accumulation of transcriptional coactivator TAZ and enhances invasiveness of human cancer cells. J Biol Chem, 297(1), 100803. 10.1016/j.jbc.2021.100803

Renier, N., Wu, Z., Simon David J., Yang, J., Ariel, P., & Tessier-Lavigne, M. (2014). iDISCO: A Simple, Rapid Method to Immunolabel Large Tissue Samples for Volume Imaging. Cell, 159(4), 896–910. 10.1016/j.cell.2014.10.010

Rinon, A., Molchadsky, A., Nathan, E., Yovel, G., Rotter, V., Sarig, R., & Tzahor, E. (2011). p53 coordinates cranial neural crest cell growth and epithelial-mesenchymal transition/delamination processes. Development, 138(9), 1827–1838. 10.1242/dev.053645

Stallings, E. B., Isenburg, J. L., Rutkowski, R. E., Kirby, R. S., Nembhard, W. N., Sandidge, T.,…National Birth Defects Prevention, N. (2024). National population-based estimates for major birth defects, 2016–2020. Birth Defects Res, 116(1), e2301. 10.1002/bdr2.2301

Stottmann, R. W., Choi, M., Mishina, Y., Meyers, E. N., & Klingensmith, J. (2004). BMP receptor IA is required in mammalian neural crest cells for development of the cardiac outflow tract and ventricular myocardium. Development, 131(9), 2205–2218. 10.1242/dev.01086

Tai, C.-J., Hsu, C.-H., Shen, S.-C., Lee, W.-R., & Jiang, M.-C. (2010). Cellular apoptosis susceptibility (CSE1L/CAS) protein in cancer metastasis and chemotherapeutic drug-induced apoptosis. Journal of Experimental & Clinical Cancer Research, 29(1), 110. 10.1186/1756-9966-29-110

Tanaka, T., Ohkubo, S., Tatsuno, I., & Prives, C. (2007). hCAS/CSE1L associates with chromatin and regulates expression of select p53 target genes. Cell, 130(4), 638–650. doi:10.1016/j.cell.2007.08.001

Tang, W., Martik, M. L., Li, Y., & Bronner, M. E. (2019). Cardiac neural crest contributes to cardiomyocytes in amniotes and heart regeneration in zebrafish. Elife, 8. 10.7554/eLife.47929

Wang, J., Xiao, Y., Hsu, C. W., Martinez-Traverso, I. M., Zhang, M., Bai, Y.,… Martin, J. F. (2016). Yap and Taz play a crucial role in neural crest-derived craniofacial development. Development, 143(3), 504–515. 10.1242/dev.126920

Zhang, X., Zhang, X., Mao, T., Xu, H., Cui, J., Lin, H., & Wang, L. (2021). CSE1L, as a novel prognostic marker, promotes pancreatic cancer proliferation by regulating the AKT/mTOR signaling pathway. J Cancer, 12(10), 2797–2806. 10.7150/jca.54482

