## Supplemental Figure 1 for "*Cse1l* Regulates Neural Crest Cell Survival and is Critical for Craniofacial and Cardiac Development"

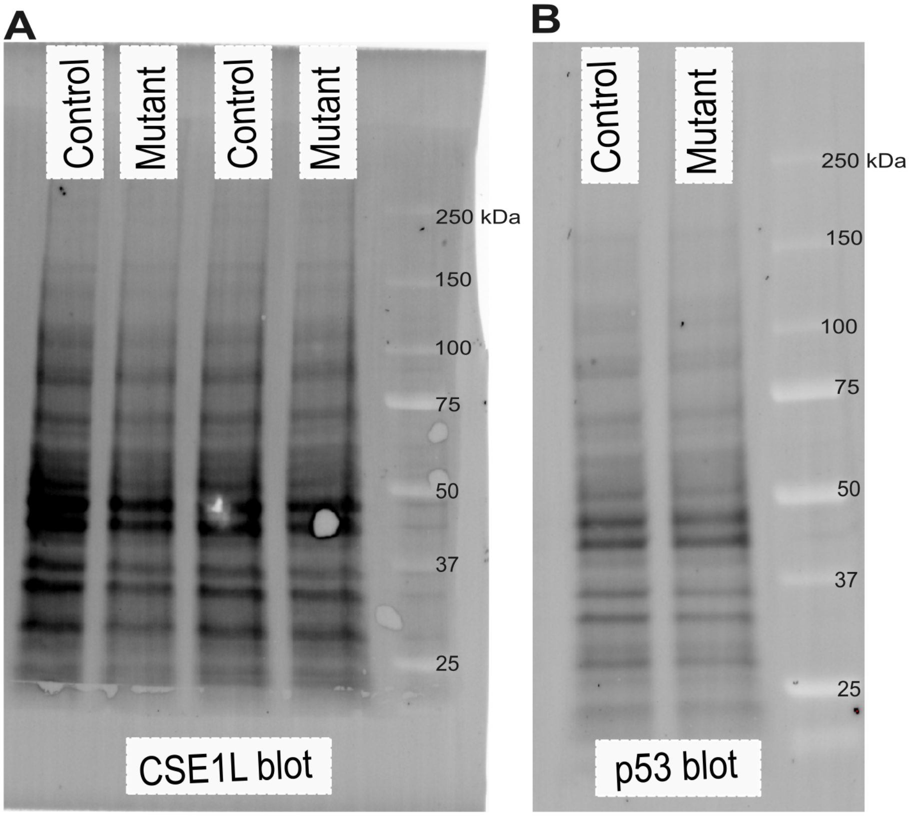


**Fig. S1.** Total protein blots used for normalization of band intensities for CSE1L (A) and p53 (B) blots. The stain-free blots were imaged prior to immunostaining were quantified for normalization for Fig. 1C for CSE1L and Fig. 7D for p53.
